# Effects of host species and environmental factors on the prevalence of *Batrachochytrium dendrobatidis* in northern Europe

**DOI:** 10.1101/349787

**Authors:** Simon Kärvemo, Sara Meurling, David Berger, Jacob Höglund, Anssi Laurila

## Abstract

The fungal pathogen *Batrachochytrium dendrobatidis* (*Bd*) is a major threat to amphibian populations. Here we asked if the prevalence of *Bd* differs between amphibian species and whether it is related to local environmental factors in breeding habitats as well as landscape variables measured at three scales (500, 2000 and 5000 m radius) in southernmost Sweden. We sampled 947 anurans from six species in 31 ponds. Canopy cover, pond perimeter, pH and temperature were treated as local scale pond characteristics. Number of surrounding ponds, area of arable land, area of mature forest and number of resident people were treated as landscape variables. *Bufo bufo* and *Rana temporaria* had a prevalence of 0.5-1.0% which differed strongly from the other four species (*Bombina bombina, Bufotes variabilis, Epidalea calamita, Rana arvalis*) showing 13-64% prevalence. *Bd* prevalence in these four species was higher in ponds with higher pH, surrounded by a landscape with less mature forest and few wetlands. Our results show that the infection dynamics of *Bd* are complex and depend on local pond characteristics, host community composition and the spatial scale under investigation. Information on environmental factors associated with *Bd* and species differences in susceptibility may mitigate further spread of the disease through public information and guide conservational action plans, especially for the most threatened species.

## Introduction

The disease dynamics of pathogens that infect multiple host species is likely to differ from that of pathogens restricted to a single host [1]. Understanding the dynamics of multiple-host pathogens is therefore especially important for conservation. For example, infection risk may be higher for less abundant species and endangered species my consequently face higher risk of decline or even extinctions [2]. Moreover, environmental factors that influence the dynamics of wildlife diseases are often unknown and vary across spatial scales [3, 4]. The occurrence of disease often varies across the range of hosts, and identifying habitats and environmental factors with high disease risk can be pivotal for mitigating the disease [5, 6]. At local spatial scales, the occurrence of infections can be affected by several factors including habitat quality [7], host population density [8], host migration [9] and interactions among host species [10]. At a regional scale, the risk of infections may additionally be affected by factors such as landscape characteristics and microclimatic conditions [11, 12]. Anthropogenic impact due to agricultural intensification, urbanization, and disease transport can also affect the infection risk at different spatial scales [3, 13].

Amphibians are the most threatened vertebrate taxon and in worldwide decline [14, 15]. One of the most serious threats to amphibian populations is the chytrid fungus *Batrachochytrium dendrobatidis* (*Bd*). This pathogen, detected in ca. 700 amphibian species [16], has caused mass mortality and population declines all over the world [14, 15, 17], including extinction of more than a hundred species [18]. *Bd* infection can result in chytridiomycosis, a disease affecting cells in the epidermis and the outer keratinized layers, disrupting the transport of water, oxygen and salts. *Bd* does not occur uniformly in the landscape as ponds and wetlands holding amphibians typically differ in their infection status [e.g. 19–21]. *Bd* prevalence differs among amphibian species which have been explained by differences in e.g. habitat preference [19, 22], immune response [23] and temperature in the breeding habitat [24]. However, although the disease is coined “the worst infectious disease ever recorded among vertebrates in terms of the number of species impacted, and its propensity to drive them to extinction” [25], there are significant knowledge gaps on differences in resistance among amphibian species and dispersal of *Bd* across landscapes.

To understand the transmission of *Bd* across species and the variation in occurrence in natural habitats, more knowledge of the factors affecting the occurrence of the fungus at different spatial scales in the landscape is required. While temperature and precipitation appear the most important environmental factors affecting the occurrence of *Bd* at large geographical scales [16, 26], factors affecting *Bd* at smaller scales seem to be more multifactorial and ambiguous. The optimal temperature for *Bd* growth in the laboratory is 17-23 °C [27], but the fungus may maintain high fitness at temperatures as low as 10 °C [28]. Several studies have shown that microclimatic habitat variation is important for *Bd* occurrence. In tropical studies, cool forests ponds or forest-dwelling individuals have a higher prevalence than warmer open-canopy ponds or individuals from open areas [6, 29, 30, but see 31]. As dry conditions reduce survival of *Bd* [32], amphibians in large permanent ponds and streams face a higher infection risk than amphibians in temporary ponds with higher desiccation risk [19, 30, 31, 33]. Similarly, a high number of surrounding ponds is associated with increased infection risk [30, 34]. In Europe, the amount of agricultural areas has been found to increase the risk [33], and in both subtropical and temperate zones the extent of surrounding forest, as well as amphibian species richness increase the risk of *Bd* [20, 26, 30, 35]. Additionally, urbanization and human activities can increase *Bd* infection risk, as humans may directly or indirectly act as vectors, or because high environmental stress due to pollution may affect host immune system [20, 33, 36, but see 37–38].

Few studies have investigated factors affecting *Bd* infection risk in cool temperate or boreal climates with relatively low summer temperatures [but see 39–42] where climate and, consequently, disease dynamics differ from those in warmer countries. In Sweden, *Bd* was first discovered in 2010 [43], followed by records on several amphibian species in southernmost Sweden and the Stockholm area, providing the northernmost records of *Bd* in Europe (S. Meurling et al., in prep.). However, Garner et al. [44] did not find *Bd* in 197 Swedish samples of museum specimens collected between 1994 and 2004, raising the possibility that *Bd* has colonized Sweden relatively recently. Currently, the environmental factors affecting the occurrence of *Bd* in northern Europe and at higher latitudes remain largely unexplored [33].

The aim of this study was to investigate the environmental factors influencing the distribution of *Bd* across different landscape scales in southern Scandinavia. To address this issue, we sampled adult amphibians from breeding aggregations in 31 wetlands in southernmost Sweden, We analysed the prevalence (proportion of infected individuals in each pond) of *Bd* while considering the effects of different species in the models, and focused on three main questions: (i) Which environmental factors influence the prevalence of Bd? (ii) Are local pond characteristics or landscape factors more important? (iii) At which scales are the landscape factors most influential?

## Materials and Methods

In total, we sampled 947 anurans from six species in 31 ponds in the province of Scania, southern Sweden, in March-May 2015 and April 2016 (Fig 1, Table 1). The study area is mainly comprised of arable lands scattered with mixed woodlands. Two of the study species, the common toad *Bufo bufo* and the common frog *Rana temporaria* are common and widespread across Sweden, including Scania. However, as only 5 of 341 (1%) common toads and 1 of 197 (0.5%) of common frogs were infested, these two species were excluded from the main analyses. The remaining four species, the moor frog *R arvalis,* the fire-bellied toad *Bombina bombina,* the green toad *Bufotes variabilis*, and the natterjack toad *Epidalea calamita* are all protected under the European Habitats Directive [45]. Of these, *R. arvalis* is a common and widespread species whereas *B. bombina, B. variabilis* and *E. calamita* are uncommon and local species in Sweden, and their distributions are limited to the southern part of the country. *B. variablis* and *E. calamita* are listed as vulnerable (VU) in the Swedish red list, *B. bombina* was removed from the list in 2015, whereas *R. arvalis* has not been not included in the red list in Sweden [46].

**Figure 1.**
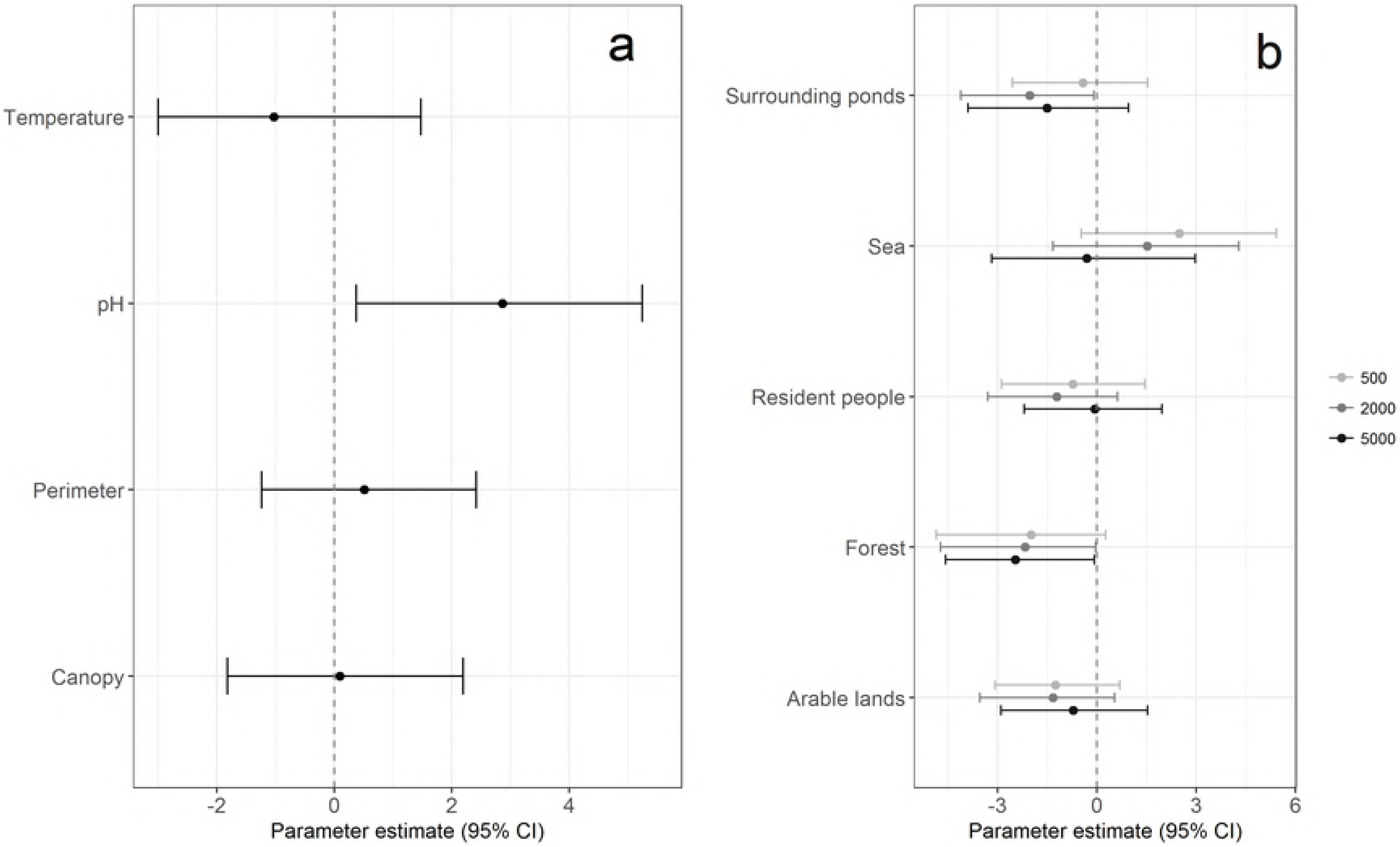
Site map. Location and prevalence (black denotes the proportion of recorded individuals with *Bd*) of the 31 studied ponds in southern Sweden. Sites with X are exclusively *Bufo bufo* and *Rana temporaria* with very low prevalence. They were not included in the main models.

The study ponds were selected based on earlier knowledge and by locating possible habitats from maps. Twenty-three ponds were sampled in 2015 and thirteen in 2016, five ponds being sampled in both years. For *Bd* sampling, we exclusively captured adults from breeding aggregations by hand or with nets, the mean number of individuals sampled being 30.5 per pond (median 25, min. 5, max. 97). We used a standard swabbing protocol with 25 strokes per individual (Brem et al. 2007). The swabs were preserved in alcohol in 2015 [47] and in Dryswab™ (MWE MW110) in 2016. No significant differences in prevalence of *Bd* were found between the years (2015=0.168, 2016= 0.178, χ^2^ = 0.00, *p*=1.00).

### Molecular analyses

DNA was extracted from swabs using the Qiagen DNeasy Blood and Tissue kit (Qiagen, ref. 69506) following the standard protocol with modifications suggested by Kosch and Summers [48]. Presence of *Bd* was assessed by amplifying the internal transcribed spacer (ITS)-5.8S rRNA region [49]. The 25μl reactions containing 12.5μl 2X Taqman Master Mix (Applied Biosystem, ref. 4318157), 2.25 μl 10μM each of forward and reverse primers, 0.625 μl 10μM MGB probe and 5μl of DNA (diluted x10 in water) were run. Each sample was run in triplicate. To avoid false negatives caused by inhibitors an exogenous internal positive control (IPC; [49]) was added to one of each triplicate (1μl 10XExo IPC master mix and 0.5μl 50XExo IPC DNA to each sample) (VICTIM dye, Applied Biosystems ref. 4304662).

**Table 1.** Infected individuals and site information. Number of individuals for each of the species included in the study, divided in *Bd*-positives, *Bd*-negatives and the prevalence. In addition, sampling periods, mean pH, and mean temperature for each study site.

| Species | Pos | Neg | Prevalence | Prevalence | Sampling | Sampling | Mean | Mean |
| --- | --- | --- | --- | --- | --- | --- | --- | --- |
|  |  |  |  | 95% CI | 2015 | 2016 | pH | temp |
| *Bombina bombina* | 10 | 19 | 34% | 18-54% | 08/5-17/5 | - | 6.92 | 12.80 |
| *Bufo bufo* | 5 | 336 | 1% | 0.5-3.3% | 27/3-11/4 | 04/4-17/4 | 7.13 | 13.01 |
| *Bufotes variabilis* | 86 | 48 | 64% | 55-72% | 01/5-21/5 | 02/5-09/5 | 7.82 | 12.77 |
| *Epidalea calamita* | 33 | 50 | 40% | 29-51% | 01/5-11/5 | 10/5-10/5 | 7.60 | 13.75 |
| *Rana arvalis* | 21 | 142 | 13% | 8-19% | 27/3-14/4 | 02/4-12/4 | 6.88 | 12.81 |
| *Rana temporaria* | 1 | 196 | 1% | 0.0-3.0% | 25/3-11/4 | 07/4-17/4 | 6.89 | 12.79 |

The qPCR assays were run on a Biorad CFX96 using amplification conditions in Boyle et al. [49] with standards of 0.1, 1, 10 and 100 genomic equivalents (GE) (2016 samples) or a positive control of approximately 1-10 GE (2015 samples). An individual was scored as *Bd* positive if one or more of the triplicate samples exhibited a positive signal (i.e. an exponential amplification curve). If the IPC showed signs of inhibition, negative samples were rerun once before being assigned as unscoreable.

### Pond variables

Four local habitat characters of each pond were evaluated: *Perimeter*, *Temperature, pH* and *Southern canopy cover*. Pond perimeter was highly correlated with pond size and chosen in the model as vegetated shallow shores are important breeding aggregation habitats of amphibians [50]. *Perimeter* was extracted with ArcMap (ArcGIS, ESRI, Redland, CA, USA) using topographic maps (Swedish National Land Survey). Ponds not visible from satellite maps were measured from polygons drawn in Google Earth and transformed to perimeter in Earth Point (www.earthpoint.us/Shapes.aspx). In cases where amphibians inhabited large lakes, *Perimeter* was based on the surface covering 500 m radius from the breeding area. In October 2017, *Temperature* (mean of three recordings per pond), *pH* (single recording measured with ThermoFisher© Orion 131S pH-meter) and *Canopy cover* (canopy along southern shore; hereafter *Canopy*) were recorded for each pond. We used *Canopy* along the southern half of the shore as these trees form effective shadow on the pond. It was estimated visually by the same person into 10% categories and strongly correlated with total percentage of surrounding large trees within 20 m from the shore (r= 0.87). Ponds with more canopy cover have lower and less variable water temperatures than open-canopy ponds [51].

### Landscape variables

We extracted a correlation matrix including all local and landscape variables and a single variable from variable pairs with a correlation coefficient >0.7 was retained in the models in order to avoid co-linearity. Consequently, we excluded three landscape variables (*total perimeter of surrounding ponds, total length of roads* and *marshland area*) from the seven initial variables. The four remaining variables *number of surrounding ponds* (hereafter *Surrounding ponds*), *area of arable lands* (crops and fruit farms, hereafter *Arable lands*), *area of mature forest* (timber volume *>300 m^3^ha^−1^*, hereafter *Mature forest*), and *number of resident people* (hereafter *Resident people*) were determined in three nested circular buffer zones with width of 500, 2000, 5000 m around the perimeter of the focal ponds, using the function “spatial join” in ArcMap. The buffer sizes were selected based on general maximum movement distances of amphibians [53]. An additional binary variable - *presence of sea* (hereafter Sea) - was included in the models to account for the negative relationship with land area.

The variables *Surrounding ponds*, *Arable lands*, and *Sea* were extracted using topographic vector maps from the Swedish National Land Survey. *Arable lands* were converted to raster format and quantified with 10 × 10 m resolution. *Mature forest* was quantified using kNN-raster data obtained from the Swedish Forest Agency [54] originally at 25 × 25 m resolution and aggregated to 50 × 50 m by averaging, due to low accuracy at the original scale [55]. *Resident people* was summed from a geographical point-layer data created by the Swedish Bureau of Statistics. To avoid confounding effects of the differences in buffer area around ponds of different size, and as some of the ponds were very small, all landscape variables were corrected by a factor calculated by dividing each buffer area (a full circle with a radius corresponding the buffer zone size) by the smallest buffer area at each landscape scale [56]. The landscape data were processed in ArcMap (ArcGIS 10.4, ESRI, Redland, CA, USA). All continuous variables were standardized to a mean of zero using the scale function in R [57].

### Statistical analyses

Effects of pond and landscape variables as well as species identity were analysed with *Bd* prevalence as the binomially distributed response variable in mixed effects models with *Site* specified as a random effect to attain the correct level of replication for the fixed effects. Due to non-convergence in some of the models using traditional maximum likelihood, we ran Bayesian linear mixed effect models with Markov-chain Monte Carlo simulations available in the package MCMCglmm [58]. We used variance expanded priors for the random effects to improve sampling properties of the MCMC chain [59]. We ran the models with residual variance fixed to 1 (as standard for binomial response models) and a flat prior on the probability scale for the fixed effects, recommended when the number of observations in some cells are low ([59]: as is the case for *Bd* prevalence at some sites and in some species) and the data show near complete separation [60]. Models ran for 2.100.000 iterations, preceded by 100.000 “burn-in” iterations that were discarded. We saved every 2000th iteration, which resulted in 1000 stored and uncorrelated (all autocorrelations <0.05) posterior estimates of model parameters upon which we based our Bayesian p-values and 95% credible intervals. While we focus on the results from these more robust Bayesian models, we note that the maximum likelihood analyses gave the same qualitative results.

To test for differences in prevalence among species, we first ran an initial model including all six different amphibians, but excluding the environmental factors. To test the effects of the environmental factors we ran the model excluding the two species (*B. bufo* and *R. temporaria*) with essentially zero incidence of (and variance in) *Bd*-prevalence. The set of explanatory variables for analyses of local environmental factors were *Perimeter*, *Temperature*, *pH*, and *Canopy*. The set of explanatory variables for estimating the influence of the landscape factors were *Surrounding ponds*, *Mature forest*, *Arable lands* and *Resident people*. The three nested distance scales (500, 2000, 5000 m) were not independent of each other, and separate analyses were conducted at each scale with the same response variable. Consequently, a total of four main analyses were conducted.

We compared model weights of the MCMC models with the function model.sel from the MuMIn package implemented in R (Bartoń 2018) [61] to estimate the most important spatial scale.

## Results

Overall, *Bd* was detected in 14 (45%) of the 31 surveyed ponds, and 16% of the sampled individuals were infected by the chytrid. For the four amphibian species included in the main models, 11 of the 23 (48%) surveyed ponds, and 40% of the individuals, were infected. All species had at least one individual infected, but prevalence differed strongly among the species (Table 1). *Bufo bufo* and *R. temporaria* had much lower prevalence than the other species (Table 1 and S1) and were excluded from the following analyses. In the remaining four species prevalence varied from 13 (*R. arvalis*) to 64% (*B. variabilis*, Table 1), however, *Species* was significant in only one of the models (2000 m) when included as a factor in the analyses of environmental factors. The infected ponds were widespread across the study area (Fig 1), and prevalence was not spatially correlated for all ponds (Moran’s I= −0.058, p=0.192), or for the subset of ponds included in the analyses (Moran’s I= −0.059, p=0.627).

Among pond variables, *pH* increased the prevalence of *Bd* at values over >6.5 (Figs 2a and 3a). *Perimeter*, *Temperature* and *Canopy* had no or weak effects. *Surrounding ponds* in the landscape had a negative effect at the 2000 m spatial scale with higher prevalence (>50%) with less than 10 surrounding ponds (Figs 2b and 3b) whereas *Mature forest* had significant negative effects at the two largest scales (Fig 2b). All positive sites had less than 200 ha mature forest within 5000 m radius (Fig 3c).

**Figure 2.**
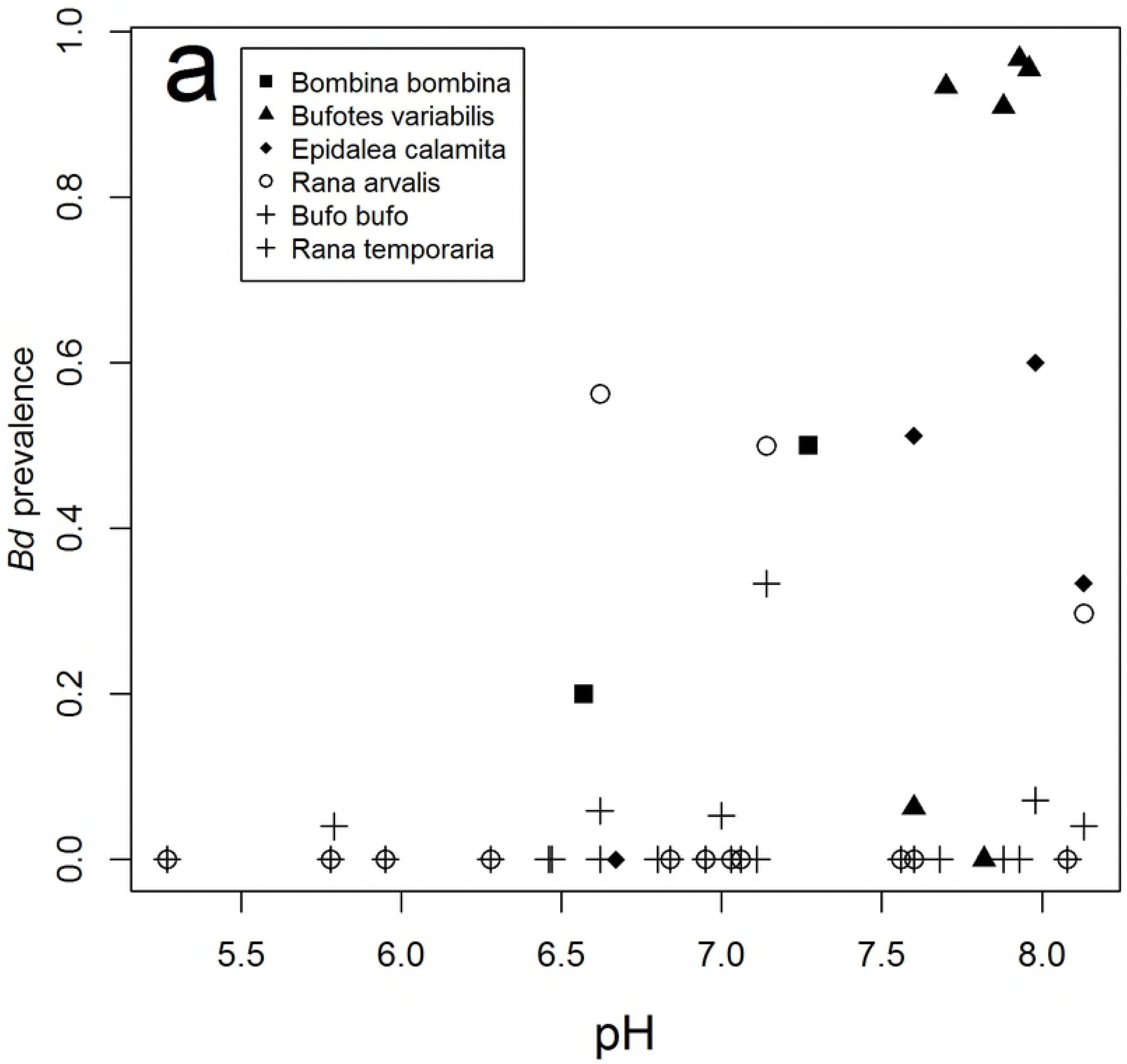
Effects on *Bd*-prevalence in the four-species Bayesian mixed-effects models. (a) Local factors (pond characteristics) and (b) environmental landscape factors. In (b) different coloured bars represent estimates at different spatial scales. Points represent posterior modes and error bars represent 95% Bayesian credible intervals.

**Figure 3.**
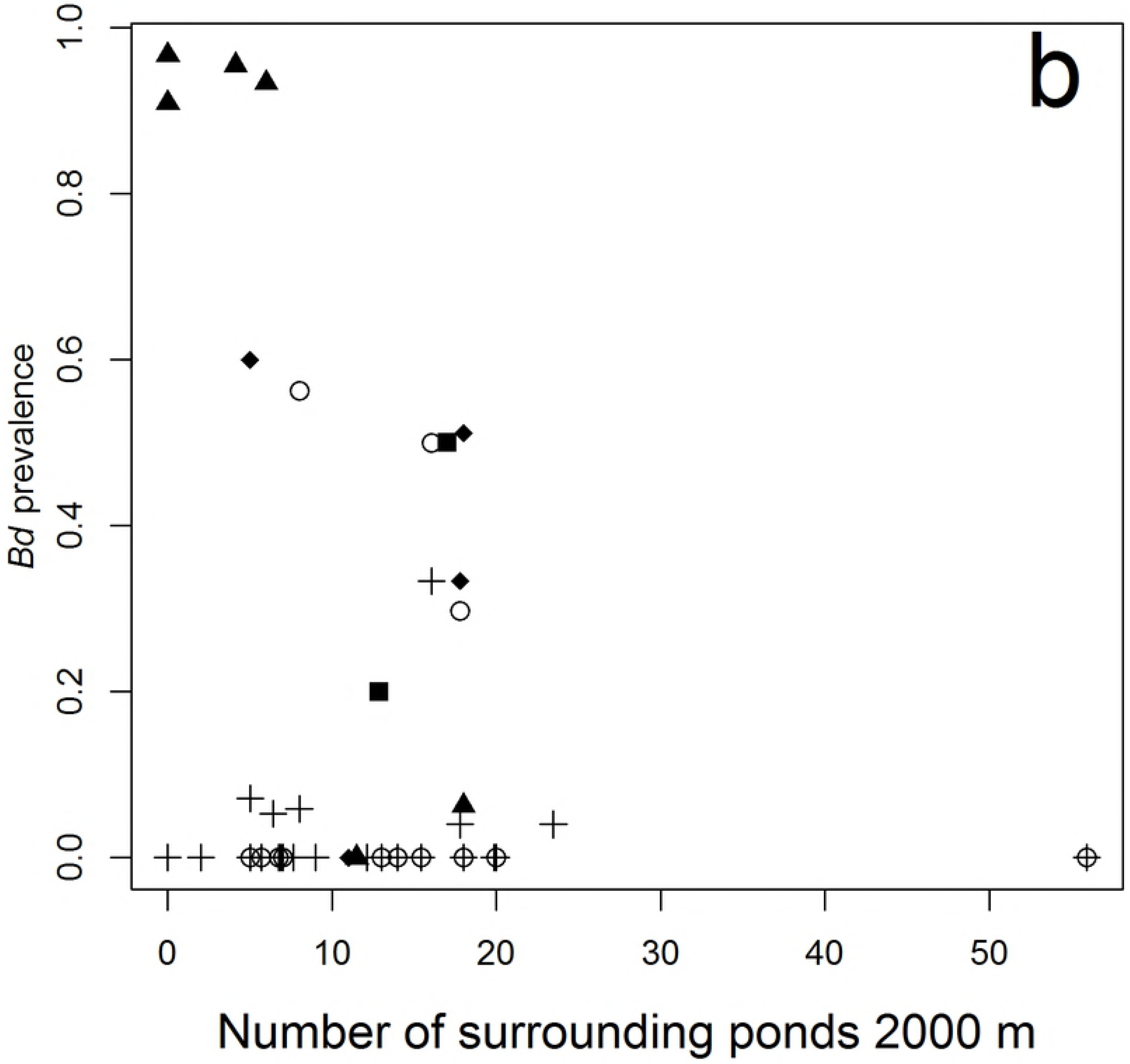
Raw data on mean estimates per site/species of prevalence and important environmental predictors from the main models. (a) Pond pH, (b) number of surrounding ponds within 2000 m, and (c) hectares of mature forests within 5000 m. The cross symbol (+) denotes species not included in the main analyses.

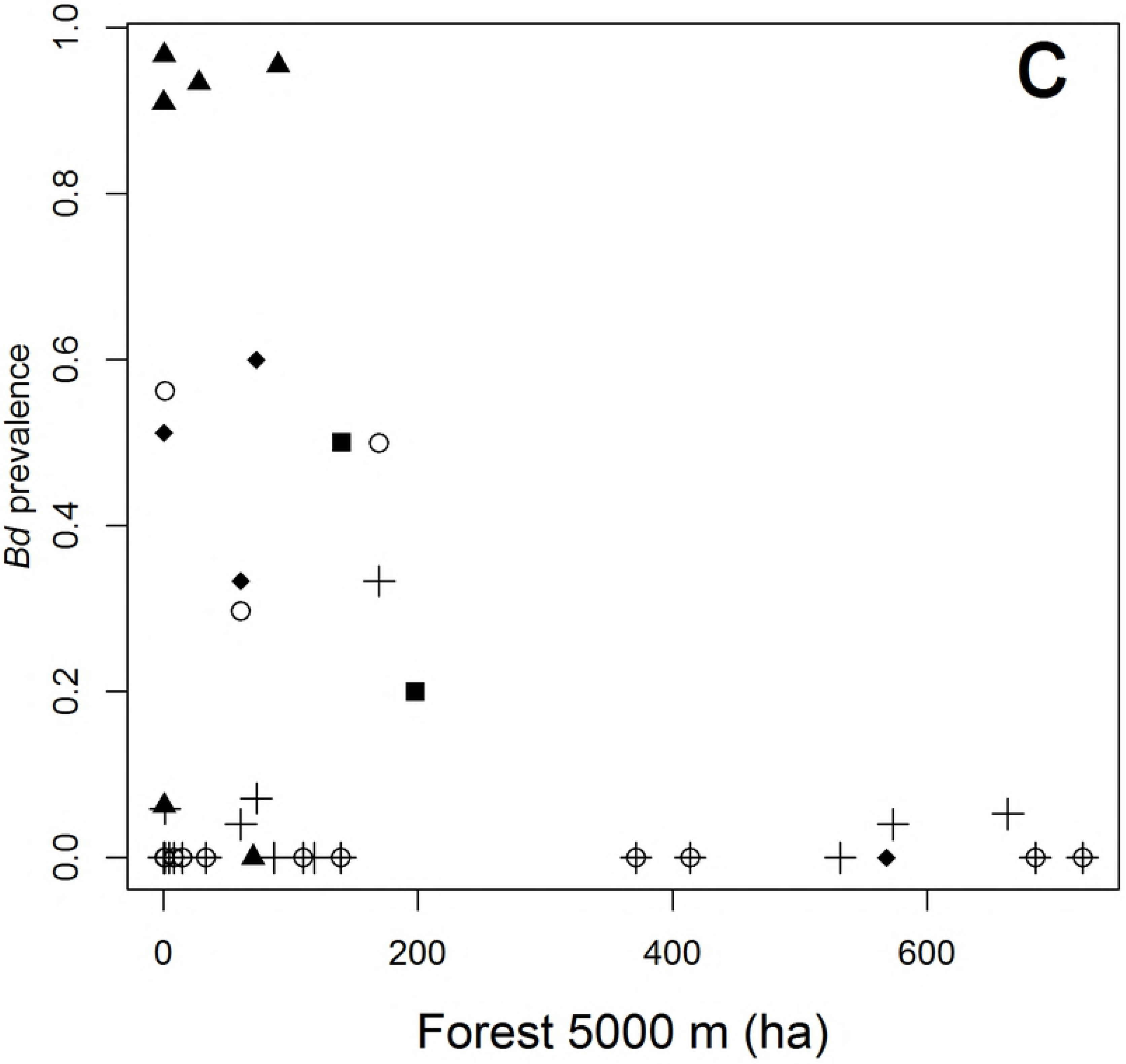

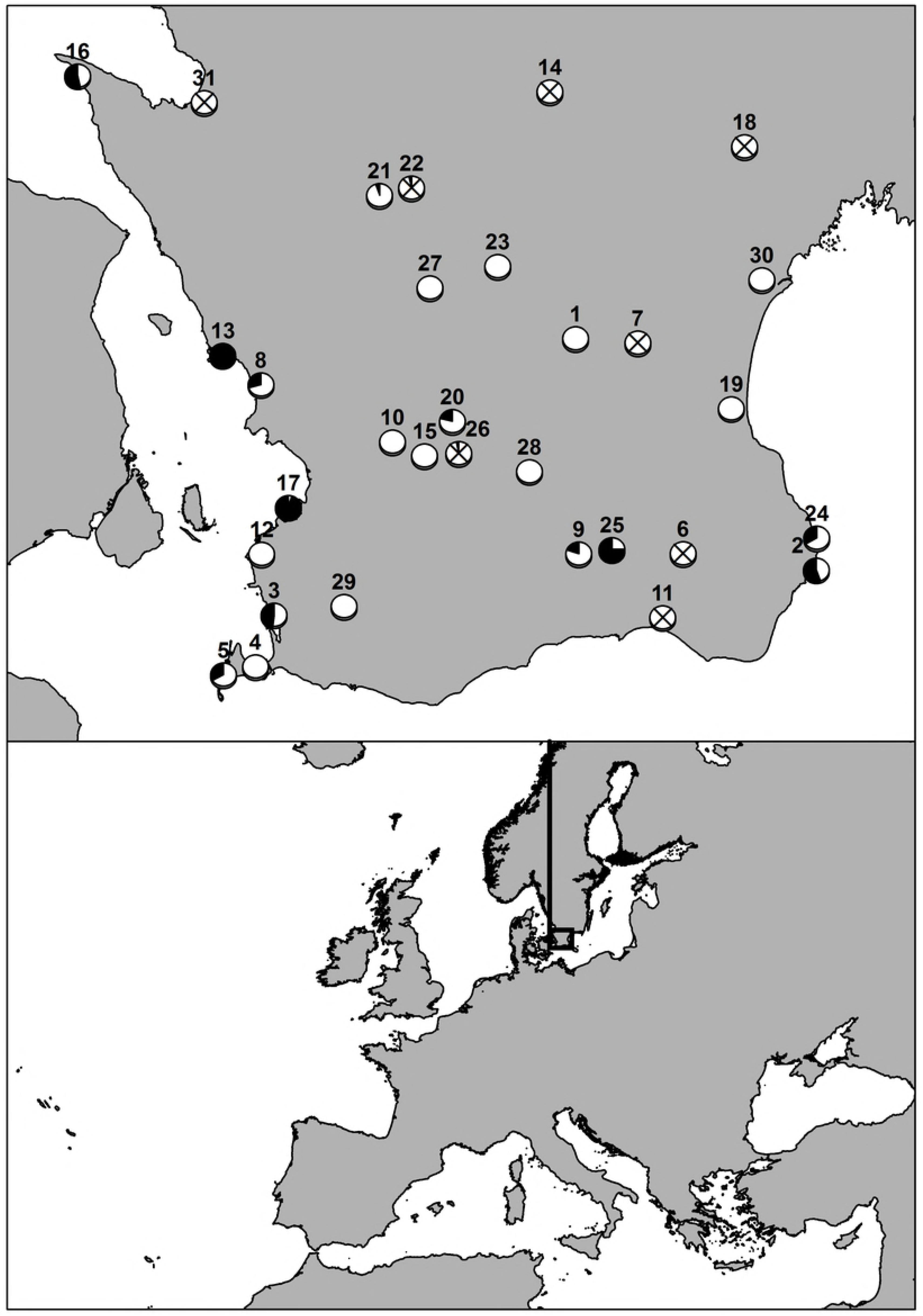

Based on the weighting of the MCMC models, the local factors explained more variation of *Bd* prevalence than the landscape factors. The environmental factors within a 5000 m landscape explained more variation in comparison to the 500 m and 2000 m scale (Table 2).

**Table 2.** Explained variation of the models. DIC (Deviance Information Criterion), BIC (Bayesian Information Criterion) and model weights of local (pond characteristics) and landscape factors generated from the MCMC models.

| Model | DIC | Model weight | BIC | Model weight |
| --- | --- | --- | --- | --- |
| Local factors | 279.0 | 0.349 | 293.9 | 0.911 |
| Landscape 500 m | 280.7 | 0.153 | 301.0 | 0.014 |
| Landscape 2000 m | 281.2 | 0.112 | 301.3 | 0.022 |
| Landscape 5000 m | 278.9 | 0.381 | 300.1 | 0.040 |

## Discussion

Understanding how *Bd* dynamics interacts with environmental factors and amphibian diversity is important for mitigating the effects of the chytrid on amphibian populations. This is the first study to address how pond characters and landscape structure affect *Bd* prevalence at high temperate latitudes. We found that host community structure as well as breeding habitat characteristics and landscape factors had impact on the prevalence of *Bd*, although it is difficult to disentangle if the environmental factors are affecting *Bd* or whether our results reflect the habitat choice of susceptible amphibian species. Our results hence emphasize the complex interplay between host community structure and local habitat and landscape characteristics.

### Species differences

Species differed in prevalence with *B. bufo* and *R. temporaria* showing the lowest prevalence (Table 1). Breeding pond temperatures for these two species are commonly below 10 °C [62–63, S. Kärvemo unpublished data]. Thus, a possible explanation for the low prevalence of *B. bufo* and *R. temporaria* are the low temperatures during breeding season of these species as warmer temperature increases the growth of *Bd* [27, 64]. The other four species breed at higher temperatures from a few days to several weeks later than the two fore-mentioned species, which may contribute to their higher prevalence. Alternatively, *B. bufo* and *R. temporaria* may have better immune defence against *Bd*, in terms of e.g. major histocompatibility complex (MHC) or antimicrobial peptide (AMP) variants coding for resistance alleles [65–66].

### Local factors

Pond pH had a strong positive effect on *Bd*, prevalence increasing when pH was higher than 6.5 (Fig 3a). This is in accordance with previous experimental (Piotrowski et al. 2004) [27] and field studies [67–68] finding higher *Bd* growth with increasing pH. It is not clear why *Bd* is affected negatively by low pH. However, Chestnut et al. [68] suggested that this may be due to reduced microbial metabolism and lower organic carbon - an important nutrient for aquatic fungi - in low pH environments.

No effects of temperature were found from the MCMC analyses, even though this has been found as an important factor in earlier studies with optimal growth of *Bd* at 17-23 °C [27, 64]. This result can be caused by the fact that temperature was not recorded during *Bd* sampling in the breeding period, but in the following autumn. However, the relative differences in water temperature should be similar between spring and autumn. Moreover, the low *Bd* prevalence of *B. bufo* and *R. temporaria*, can still be caused by a temperature effects as these two species generally were sampled approximately one month earlier than the more warm adapted species. Indirect factors associated with temperatures have been shown to affect the prevalence of *Bd*, for example, local canopy cover [6, 29] and elevation [20, 36]. In our study, canopy cover did not show any effects, probably because none of the sites included had more than 65% canopy cover and only two sites had more than 50%. This may have resulted in too little variation to detect any effects. Similarly, variation in elevation was very limited in our study (mean 37.5, max. 170.9 m) and this variable was not included in the analyses after preliminary screening.

Based on DIC (Deviance Information Criterion), local factors explained most of the model variation, or similar as the landscape factor at the largest scale. However, the BIC (Bayes Information Criterion), which penalizing the number of factors in a model, showed clearly the highest explanation for local factors. Therefore, our results suggest that local factors included in the models are somewhat more important than the landscape factors.

### Landscape factors

The most consistent landscape effects across spatial scales were the negative effects of mature forest, which had effects at the two largest scales. In contrast to our results, earlier studies have shown that forests increase the risk of *Bd* for pond-breeding amphibians (e.g. Olson et al. 2013 [26]). This is especially the case in areas with warmer climate than Sweden, e.g. California (Pauza et al. 2010) [36] and Romania (Scheele et al. 2015) [30], where the mean spring temperatures are 3-8 °C warmer than in southern Sweden (www.weatherbase.com). Accordingly, the reason for the negative impact of forests on *Bd* in our study site could be that forested environments are too cool and avoided by some amphibian species (e.g. Blomquist and Hunter Jr 2009) [69]. This could be especially the case for these four species included in the analyses, as at least three of them prefer open habitats [70].

The number of surrounding ponds had a negative impact on prevalence at the 2000 m scale. These migration distances are common for pond-breeding amphibians [53]. We suggest that the negative impact of surrounding ponds may be caused by a dilution effect, induced by an increased amphibian biodiversity and density with connectivity, which may, in turn, decrease the prevalence and infection intensity of *Bd* [71, 72]. Although the differences between the scales were small, the landscape variables at the 5000 m scale explained most variation in *Bd* prevalence. This is within a feasible dispersal distance for anurans [53] and indicate interactions of environmental factors and *Bd* at this scale.

## Conclusions

Our results suggest that the importance of environmental factors on *Bd* occurrence varies across species and spatial scales. For future investigations, it is therefore highly important to choose relevant scales and avoid drawing general conclusions based on a single species. In general, local factors seem to have more influence than landscape factors. Our results suggest that in cooler climates, the effects of landscape factors on *Bd* prevalence can differ from those found at lower latitudes. Specifically, amphibians inhabiting ponds with relatively high pH may face a higher risk of *Bd* infestation, whereas individuals inhabiting landscapes with many surrounding ponds and large areas of mature forests may have a reduced risk. Importantly, many of the environmental factors in this study are modified by humans over time. Without thorough cross-scale analyses, researchers and stakeholders may misestimate the impact of human-mediated changes in the landscape on amphibians, ecosystems and overall biodiversity [72].

## Acknowledgements

We thank David Åhlén for valuable assistance in the field and the lab. *Bd* sampling was permitted by the Skåne County Board and ethical committee for animal experiment in Uppsala county (5.8.18-04423/2017). Anders Hallengren and Jenny Hall helped with sorting out the permits. Erik Ågren provided advice on how to screen for *Bd*. Mats Wirén (Malmö municipality) and Jon Loman provided help and suggestions on identifying the study ponds.

## Supporting information

**Table S1. Six species and differences in *Bd* prevalence.** Bayesian mixed-effects models of Bd-prevalence and the six species of amphibians with *Bombina bombina* as the intercept against which prevalence in the other species was tested.

**Table S2. Four species and differences *Bd* prevalence.** Bayesian mixed-effects models of Bd-prevalence and the four species of amphibians included in the main models, with *Bombina bombina* as the intercept as the intercept against which prevalence in the other species was tested.

